# Assessing the virulence of *Cryptococcus neoformans* causing meningitis in HIV infected and uninfected patients in Vietnam

**DOI:** 10.1101/189274

**Authors:** Lam Tuan Thanh, Dena L. Toffaletti, Jennifer L. Tenor, Charles Giamberardino, Gregory D. Sempowski, Yohannes Asfaw, Hai Trieu Phan, Anh Van Duong, Trinh Mai Nguyen, Guy E. Thwaites, Philip M. Ashton, Chau Van Vinh Nguyen, Stephen G. Baker, John R. Perfect, Jeremy N. Day

**Affiliations:** Oxford University Clinical Research Unit, Wellcome Trust Asia Africa Programme, Ho Chi Minh City, Vietnam; Division of Infectious Diseases, Department of Medicine and Department of Molecular Genetics and Microbiology, Duke University, North Carolina, USA; Duke Human Vaccine Institute and Regional Biocontainment Laboratory, Duke University, North Carolina, USA; Division of Laboratory Animal Resources, Duke University, North Carolina, USA; Centre for Tropical Medicine and Global Health, Nuffield Department of Medicine, University of Oxford, Oxford, UK; Hospital for Tropical Diseases, Ho Chi Minh City, Vietnam; Cambridge Institute of Therapeutic immunology and Infectious Disease, Department of Medicine, University of Cambridge, Cambridge, UK

**Keywords:** Cryptococcal meningitis, *Cryptococcus neoformans*, immunocompetent, HIV, virulence

## Abstract

We previously observed a substantial burden of cryptococcal meningitis in Vietnam atypically arising in HIV-uninfected individuals. This disease was associated with a single genotype of *Cryptococcus neoformans* (Sequence Type (ST)5), which was significantly less common in HIV-infected individuals. Aiming to compare the phenotypic characteristics of ST5 and non-ST5 C. neoformans we selected 30 representative Vietnamese isolates, compared their *in vitro* pathogenic potential and *in vivo* virulence. ST5 and non-ST5 organisms exhibited comparable characteristics with respect to *in vitro* virulence markers including melanin production, replication at 37°C, and growth in cerebrospinal fluid. However, the ST5 isolates had significantly increased variability in cellular and capsular sizing compared with non-ST5 organisms (*p*<0.001). Counter-intuitively, mice infected with ST5 isolates had significantly longer survival with lower fungal burdens at day 7 than non-ST5 isolates. Notably, ST5 isolates induced significantly greater initial inflammatory responses than non-ST5 strains, measured by TNF-α concentrations (*p*<0.001). Despite being generally less virulent in the mouse model, we hypothesize that the significant within strain variation seen in ST5 isolates in the tested phenotypes may represent an evolutionary advantage enabling adaptation to novel niches including apparently immunocompetent human hosts.

## Introduction

Cryptococcal meningitis (CM) is a life-threatening fungal infection caused principally by *Cryptococcus neoformans* and *Cryptococcus gattii* ^1–3^. Despite CM being more common in immunocompromised patients, CM can also arise in apparently immunocompetent patients. Recent figures estimate an annual global CM incidence of 223,100 cases in HIV infected patients, resulting in 181,100 deaths ^4^.

Generally, *C. gattii* is the most common cause of CM in immunocompetent patients, while *C. neoformans* is primarily responsible for disease in immunocompromised patients ^5^. When CM associated with *C. neoformans* var. *grubii* occurs in HIV-uninfected individuals, reports often describe patients with an increased susceptibility due to other underlying immunosuppressive conditions ^6,7^. In Vietnam, *C. neoformans* var. *grubii* CM in HIV-uninfected patients accounts for approximately 20% of all CM cases admitted to our hospital in Ho Chi Minh City (HCMC) ^8^. We previously reported that the majority of HIV-uninfected CM patients had no identified cause or other medical history suggestive of immunosuppression, and that >80% of HIV-uninfected CM patients were infected by a single genotype of *C. neoformans* var. *grubii* ^9^, which we later confirmed to be Sequence Type 5 (ST5).

Our clinical observations have been replicated in China, where >70% of CM cases from apparently immunocompetent individuals were infected with *C. neoformans* ^10,11^. Again these organisms were latterly identified as genotype ST5 ^12,13^. Comparably, ST5 accounted for >80% of HIV-uninfected CM patients in South Korea, although some patients from this location had potentially immunosuppressive conditions ^14^. This association between a ST and host immune phenotype could be explained by lineage-specific increases in pathogenic potential, fitness in the human host, an unidentified host immune deficit, or a combination of these three factors. Recent high-resolution genomic investigation of the population structure of clinical *C. neoformans* var. *grubii* isolates in Vietnam has verified that ST5 and ST4 are genetically distinct lineages of *C. neoformans* var. *grubii*, namely VNIa-5 and VNIa-4 ^15^.

Here, in order to explore the hypothesis that *C. neoformans* var. *grubii* ST5 have an increased pathogenic potential in comparison to non-ST5 organisms, we compared their *in vitro* virulence phenotypes and exploited an established cryptococcosis mouse model to compare their relative pathogenicity and ability to induce systemic inflammation.

## Materials and methods

### C. neoformans isolates and culture conditions

We used clinical isolates obtained at the point of diagnosis, prior to antifungal therapy, from the cerebrospinal fluid (CSF) of patients enrolled in a prospective descriptive study of HIV-uninfected patients with central nervous system (CNS) infections, and a randomized controlled trial of antifungal therapy in HIV-infected patients ^8,16^. We randomly selected 15 isolates from HIV uninfected CM patients and 15 from HIV-infected CM patients. MLST profiles (CAP59, GPD1, IGS1, LAC1, PLB1, SOD1, URA5) for all isolates were previously determined ^17^. *C. neoformans* yeasts were propagated using Yeast Peptone Dextrose (YPD) broth and incubated overnight at 30°C with agitation. Isolates and clinical information from corresponding patients are summarized in Table 1.

**TABLE 1.**
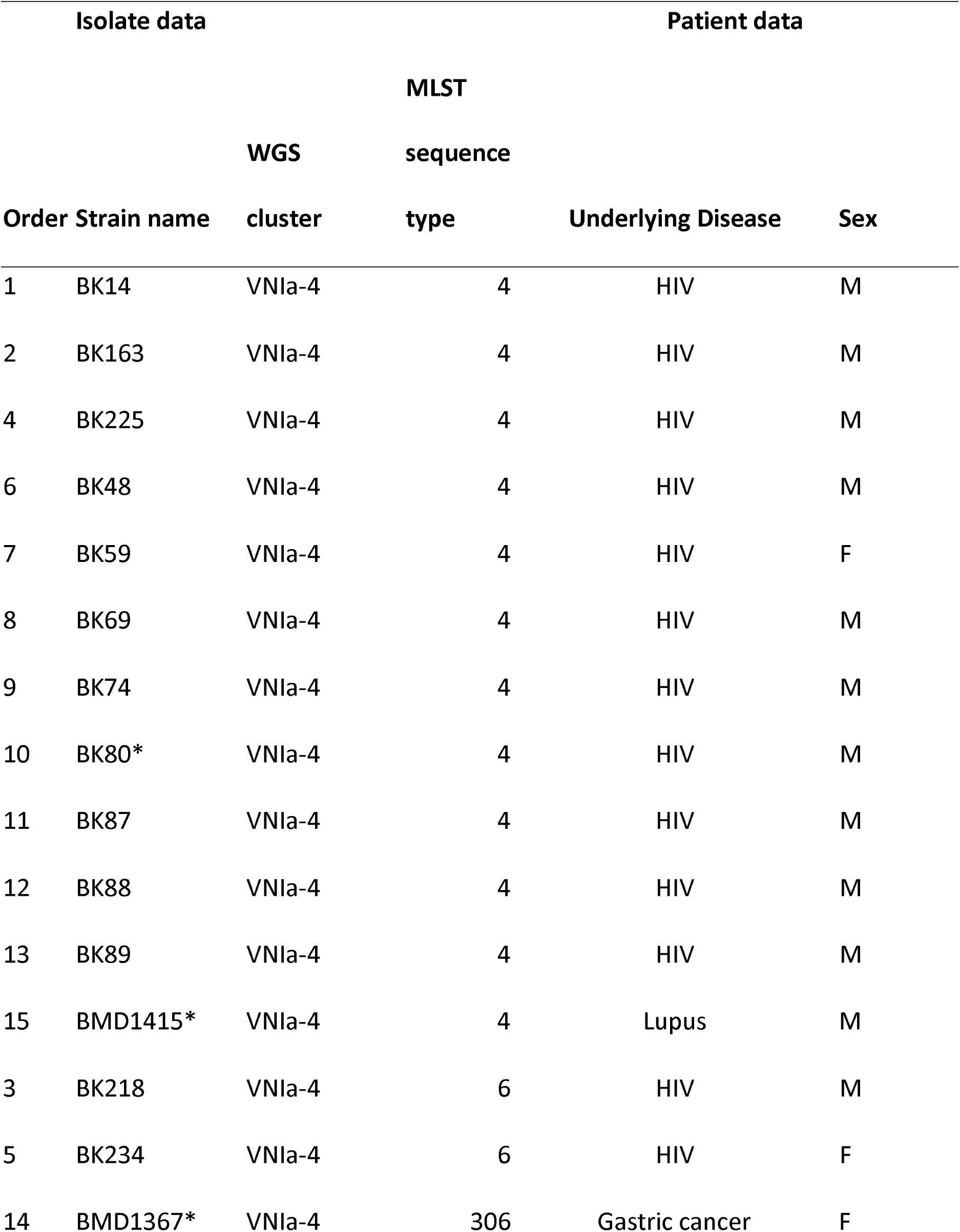
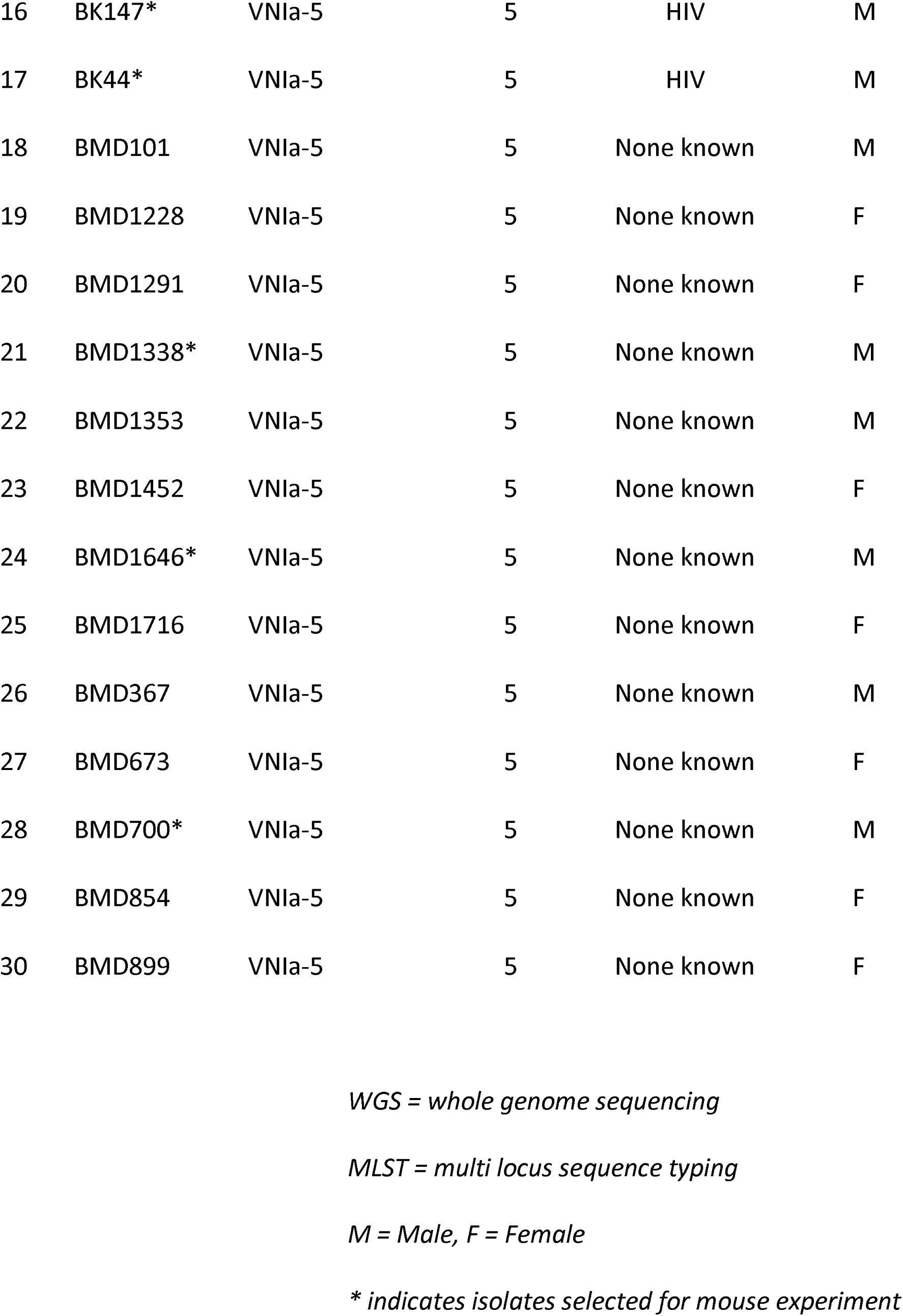
*Cryptococcus neoformans* isolate clinical source and typing (n=30).

### Growth at high temperature, in ex vivo human CSF and melanin production

Growth at high temperature and in *ex vivo* human CSF were tested as previously described ^18^ with modifications for quantitative assessment. To assess fungal growth at different temperatures, the inoculum was adjusted to 10^8^ cells/ml, serially diluted and spot-inoculated in duplicate on YPD agar in 5µl aliquots and incubated at 30°C or 37°C for 48 hours. After 48 hours, colony forming units (CFU) were counted and recorded in CFU/ml.

For the *ex vivo* CSF growth assay, baseline pre-antifungal treatment CSF supernatant from random de-identified HIV-infected patients enrolled into an antifungal therapy trial was pooled, filtered, and stored at -80°C until use. 10µl of 10^8^ cells/ml yeast suspension was inoculated into 90µl of pooled CSF and incubated at 37°C with 5%CO_2_. Inoculated CSF was serially diluted and spotted on YPD agar at days 1 and 3 post-inoculation. All experiments were repeated in triplicate. The H99-derived mutant *Δena1*, which lacks a cation-ATPase-transporter resulting in decreased viability in human CSF and macrophages, was used as a negative control for the *ex vivo* CSF assay ^19^. H99 was included as a reference in all experiments. Data were standardized by expressing the results as a ratio of the CFU/ml of the test isolate to the CFU/ml of H99. Melanin production was assessed by plating 5μl of 10^6^cells/mL cell suspension on L-DOPA agar containing 1g/L L-asparagine, 1g/L glucose, 3g/L KH_2_PO_4_, 250mg/L MgSO_4_.7H_2_O, 1mg/L Thiamine HCl, 5μg/L Biotin, 100mg/L L-DOPA, 20g/L Bacto Agar ^20,21^. Plates were incubated in the dark at 30°C or 37°C for 3 days. Differences in colony melanization were compared visually with reference to H99 and an H99-derived mutant with diminished melaninization in L-DOPA agar from the Perfect lab.

### Extracellular urease and phospholipase activity

Extra-cellular urease production was semi-quantified using Christensen’s agar. 10μl of 10^8^ cells/ml yeast suspension was spotted on Christensen’s agar and incubated at room temperature. The time to complete plate colouration was determined using a GoPro Hero 6 camera (Gopro, USA) using the time-lapse setting set with a 1-minute interval. *C. neoformans* H99 was used as a positive control and *Candida albicans* as a negative control. Extracellular phospholipase activity was screened on egg yolk medium as previously described, with minor modifications ^22^. The egg yolk medium contained Sabouraud agar with 1M sodium chloride, 0.005M calcium chloride and 8% sterile egg yolk enrichment (Merck, USA). A 5µl aliquot of *C. neoformans* yeast suspension (10^8^ cells/ml) was spotted on egg yolk agar and incubated at 30°C for 72 hours. The diameters of the precipitation zone (D) formed around the colonies and of the respective colonies (d) were recorded after 72 hours incubation. The D/d ratio for each isolate was calculated. H99 was included for reference in each experimental batch. The final result for each isolate was expressed as the ratio between the test isolate’s D/d ratio and that of H99. All isolates were tested in triplicate for each phenotype.

### In vitro capsule and cell size measurement

To measure *in vitro* cryptococcal capsule thickness, all isolates were streaked onto capsule-inducing agar containing powdered Dulbecco Modified Eagle Medium (DMEM) [supplemented with 4.5g/L glucose, L-glutamine, sodium pyruvate], NaHCO_3_ 250mM, NaMOPS 1M, Neomycin 200mg/ml, Cefotaxime 100 mg/ml ^23^. Plates were incubated at 37°C in 5% CO_2_ until single colonies were visible. Unless otherwise specified, all reagents were purchased from Sigma-Aldrich. India ink smears from a single yeast colony were prepared on a glass slide and visualized at 100X magnification using a CX41 microscope (Olympus, Japan). Images of single microscopic yeast cells were captured using a DP71 Camera system with DP Controller software (Olympus, Japan) and processed using ImageJ (rsb.info.nih.gov/ij/). Capsular thickness was calculated by subtracting yeast cell body diameter (D_CD_, no capsule) from whole cell diameter (D_WC_, including capsule). At least 30 individual microscopic yeast cells were assessed for each isolate.

### Mouse inhalation infection model of cryptococcosis

All mouse infection experiments were conducted as previously described according to Duke University’s Institutional Animal Care and Use Committee guidelines and approvals ^24^. Six-week old female A/J mice were sedated with isoflurane and inoculated intranasally with the selected *C. neoformans* var. *grubii* isolate by dropping 25µl of yeast suspension containing 5×10^4^ cells into the nares. Eight isolates were randomly selected from the 30 for murine experiments. The isolates were five ST5 (BK147 and BK44 from HIV infected patients and BMD700, BMD1338 and BMD1646 from HIV uninfected patients) and three non-ST5 strains (BMD1415 (ST4) and BMD1367 (ST306) from HIV uninfected patients and BK80 (ST4) from an HIV infected patient). Animals were monitored daily and euthanized by CO_2_ inhalation at indicated time points (fungal burden and *in vivo* responses) or until weight loss ≥ 15% body weight was observed (virulence assay).

### Determining in vivo fungal burden

Five mice were infected with each isolate in two independent experiments for assessment of fungal burden at 7-or 14-days post-infection. All animals in each experiment set were euthanized by CO_2_ inhalation either on day 7 or day 14 post-infection. Fungal burden at each time point was assessed by excising the left superior lobe of the lung and brain and homogenizing the tissue by bead beating. Tissue homogenate was serially diluted and plated onto YPD agar supplemented with 100mg/ml ampicillin. The plates were incubated at 30°C for 48 hours and the number of *C. neoformans* colony forming units (CFU) recorded. Fungal burdens were expressed as CFU per gram of tissue (CFU/g). At each time point, additional lung lobes were also collected for determining *in vivo* histopathology and cytokine response, as described below. In addition, fungal burdens were separately determined at the point of death in animals from the survival assays described below.

### Determining in vivo histopathology

At specific time points (7 or 14 days post-infection, as described above), the right superior lung lobe from each mouse was excised and immersed in 10% formalin (replaced with 70% ethanol after 24 hours) for fixation. Fixed, uninflated lung specimens were stored at 4°C until further processing. After paraffin embedding, sliced sections were stained using the periodic acid-Schiff (PAS) or mucicarmine stains. Histopathological examination was performed by an independent pathologist blinded to infecting strain. Tissue damage was scored from 0 (no changes) to 10 (severe changes), corresponding to the severity of pathology in 4 different categories: necrosis, hemorrhage, edema and inflammation, as per the Duke Veterinary Diagnostic Laboratory (Division of Laboratory Animal Resources).

### Determining in vivo cytokine response

To assess the severity of the inflammatory responses at specific time points (day 7 and day 14 post-infection), the middle lobe from the right lung of each infected mouse was excised and homogenized by bead beating in 1ml sterile PBS/Protease inhibitor. 500µl of lung homogenate was used for cytokine profiling. Cytokines representing T-helper type 1 (Th1) (IL-12p70, TNF-α, IFN-γ), T-helper type 17 (IL-17) and T-helper type 2 (Th2) (IL-4, IL-5, IL-10) responses were measured using a customized Bio-Plex Pro™ Mouse Cytokine Th1/Th2 Assay kit (Biorad, USA) with the BioPlex 200 platform according to the manufacturer’s guidelines. Data were retrieved using BioPlex Manager Software. The upper and lower limits of quantification (ULOQ and LLOQ) were based on a standard curve. All values falling below the LLOQ were replaced with the midpoint between zero and the LLOQ. Data were standardized by lung weight and presented as picogram of cytokines per gram lung tissue (pg/g).

### In vivo virulence assay

The virulence assay was conducted independently from the day 7/day 14 experiment. Each of the 8 selected isolates was inoculated intranasally into 10 A/J mice. Mice were monitored daily until death, or loss of more than 15% body weight (impending death), at which point they were euthanized by CO_2_ inhalation, necropsied and had fungal burden in lung and brain determined.

### Statistical analysis

GraphPad Prism version 5.04 for Windows (GraphPad Software, San Diego California USA; www.graphpad.com) was used for data visualization and statistical analyses of fungal loads, cytokine profiling, capsular/cell size, and survival proportions. The Mann-Whitney U-test was used for comparing fungal load and cytokine concentrations. Kaplan-Meier survival curves and the log-rank test were used for survival analysis. Capsule/cell size was compared using Welch’s t-test. The Fligner-Killeen test of variance homogeneity for analyzing variation in capsule/cell size and Fisher’s exact test were performed using R software, version 3.2.4 (http://www.r-project.org). One way ANOVA with post hoc multiple comparison tests (Dunnett or Bonferroni) was used to compare cytokine concentrations between individual isolates.

## Results

### C. neoformans in vitro virulence

We compared the capsule size, extracellular urease, phospholipase production, melanin production, growth at high temperature, and growth in pooled human CSF between the 15 ST5 and the 15 non-ST5 *C. neoformans* isolates. We observed no significant genotype-specific differences between growth at 30°C (*p*=0.10), growth at 37°C (*p*=0.23), *ex vivo* survival in CSF after 1-day (*p*=0.72), *ex vivo* survival in CSF after 3-days of exposure (*p*=0.77), extracellular urease activity, or phospholipase activity (Figure S1). ST5 *C. neoformans* cells developed significantly thicker capsules during *in vitro* culture than non-ST5 isolates (Table 2). Individual ST5 *C. neoformans* cells were also significantly larger than non-ST5 cells (Figure 1), and we observed significantly greater variation in capsule size and cell diameter within ST5 cells than non-ST5 cells (Figure 1; *p*<0.0001, Fligner-Killeen test). Notably, a single organism (BMD1646), was clearly larger and with thicker capsules than other isolates. However, the difference in capsule thickness and cell diameter between STs remained statistically significant even when this isolate was removed from analysis.

**TABLE 2.** Variability in *in vitro* capsule thickness and cell diameter of *Cryptococcus neoformans* strains by Sequence Type (ST)

| Variable | MLST Group |  |
| --- | --- | --- |
|  | ST5 | Non-ST5 |
| **Capsule Thickness (µm)** |  |  |
| Mean* | 2.64 | 2.01 |
| 95% CI | 2.49 - 2.79 | 1.94 - 2.07 |
| Range | 0.01 - 9.56 | 0.02 - 6.16 |
| Coefficient of Variation<sup>\$</sup> | | |
| (%) | 0.63 | 0.37 |
| **Cell diameter (µm)** |  |  |
| Mean* | 11.38 | 9.69 |
| 95% CI of Mean | 10.96 - 11.79 | 9.52 - 9.87 |
| Range | 4.74 - 27.22 | 5.25 - 19.89 |
| Coefficient of Variation<sup>\$</sup> | | |
| (%) | 0.41 | 0.20 |

\* $P < 0.0001$ , *t*-test with Welch's correction.
<sup>\$</sup> $P < 0.0001$ , *Fligner-Killeen test of homogeneity of variance*.

**Figure 2.**
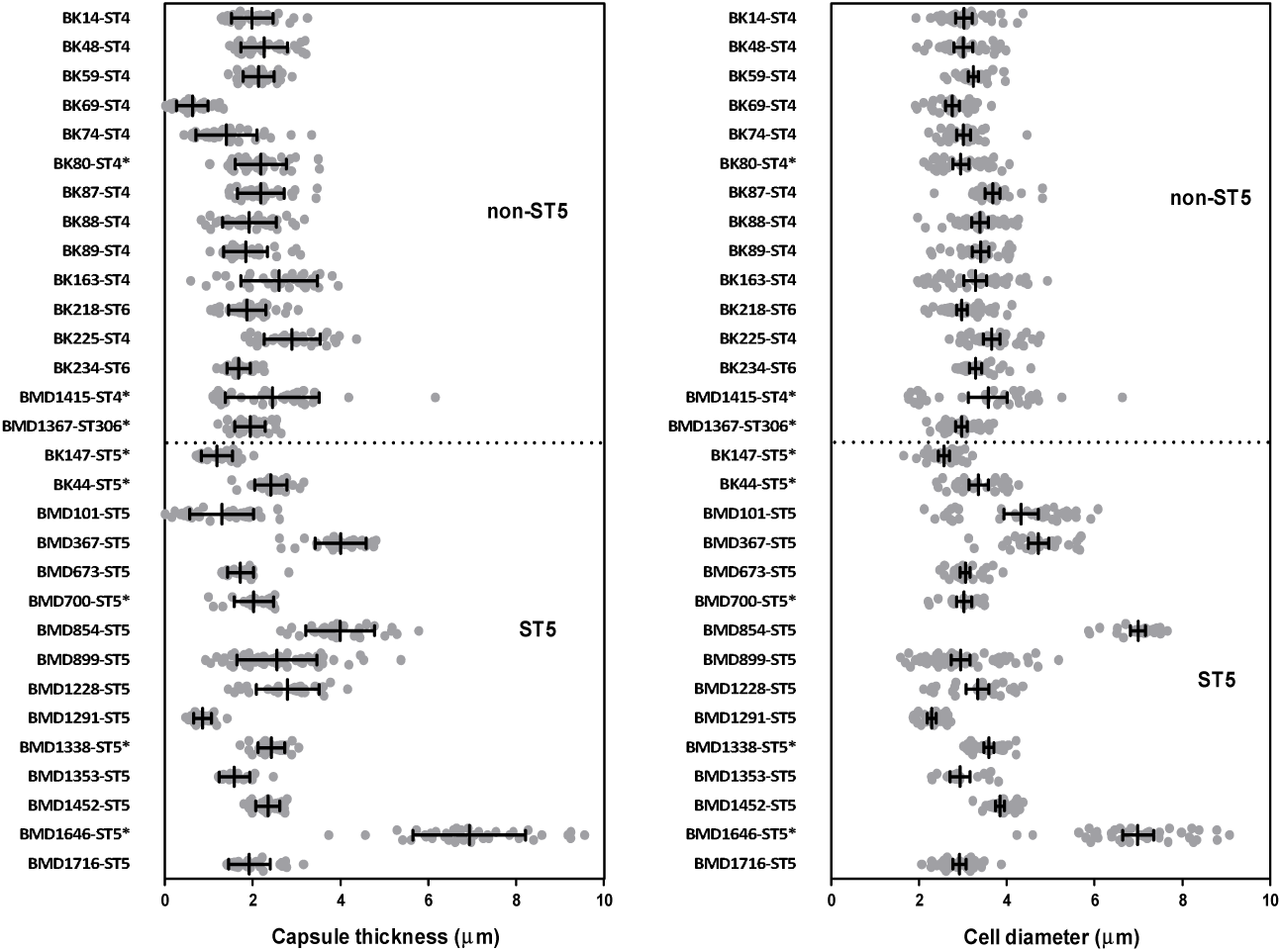
*In vitro* induced capsule thickness and cell diameter of individual *Cryptococcus neoformans* strains from Vietnam: Cells were grown on DMEM medium/5% CO_2_ and visually assessed by India ink staining. Images were taken for single cells measurement using ImageJ software. Capsule thickness is obtained by subtracting cell body diameter from total cell diameter. AFLP-VNI-γ/MLST-ST5 strains expressed higher degree of variation in both capsule size and cell diameter *in vitro*, which remains significant even when the outlier BMD1646 was removed from the analysis (*p*<0.0001 for both capsule and cell size, Fligner-Killeen test). Scattered plot represents single cells from an individual strain. Data for individual strains are presented as mean with error bars denoting standard deviation. Strains selected for experiment in mice were indicated by asterisks.

### In vivo mouse infections with ST5 and non-ST5 isolates

We hypothesized that a higher prevalence of ST5 infections in apparently healthy hosts was associated with higher virulence. Therefore, we challenged A/J with equivalent doses of representative isolates (5 ST5 and 3 non-ST5; the individual *in vitro* virulence phenotyping results for these 8 isolates are presented in supplementary Figures S2-S4). Contrary to our hypothesis, we found that mice infected with ST5 organisms had significantly longer survival times than mice infected with non-ST5 organisms (*p*<0.0001, Figure 2A). However, there was significant variability in the effect of individual organisms of the same ST on survival (Figure 2B). Specifically, two ST5 organisms (BK147 and BMD1646) were substantially attenuated in comparison to other ST5 organisms; the mice infected with these organisms survived up to 40 days. However, differences in survival times between mice infected with ST5 and non-ST5 organisms remained significant when these two isolates were removed from the analysis (*p*=0.003; log-rank test).

**Figure 2.**
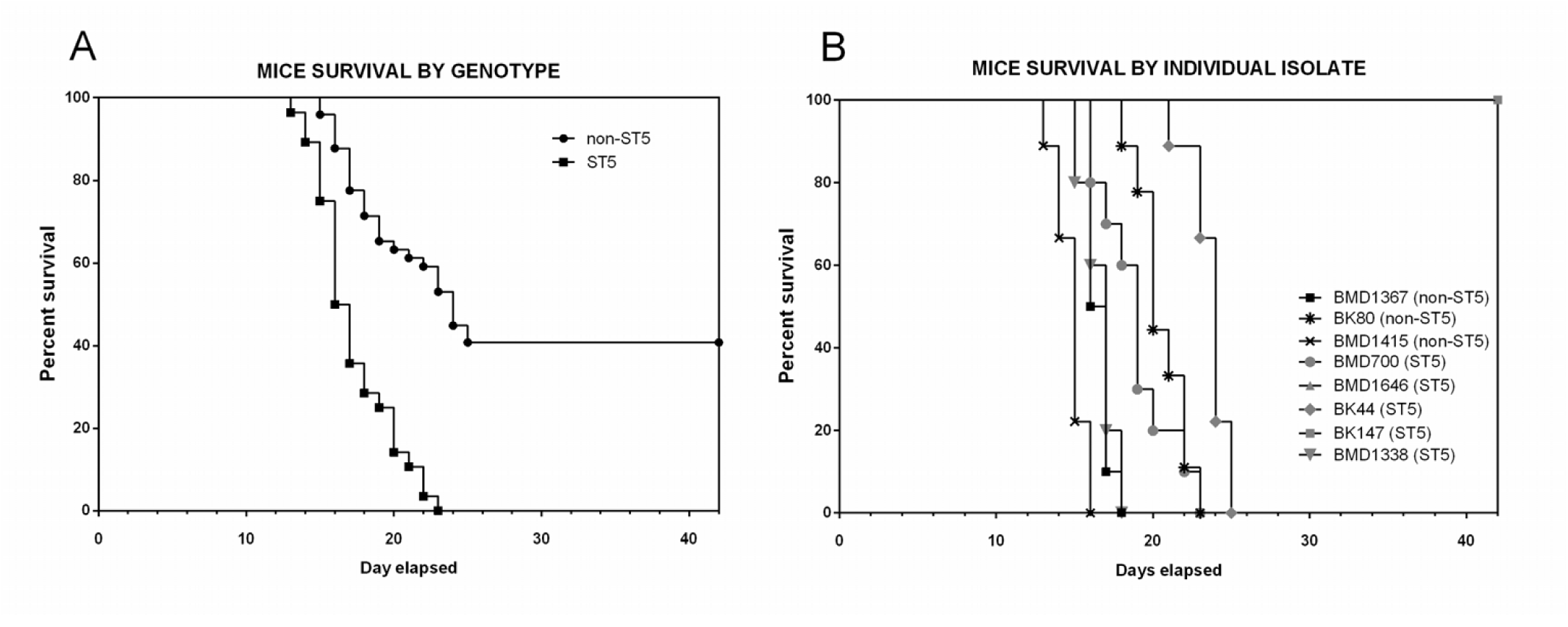
Kaplan-Meier survival curves for mice infected with either ST5 (n = 5) or non-ST5 (n= 3) *Cryptococcus neoformans* strains. 10 A/J mice were infected per strain (five ST5 strains and three non-ST5 strains). Mice were monitored daily until the point of more than 15% weight loss or visible suffering and were then sacrificed by CO_2_ inhalation. Mice infected with ST5 strains had statistically significantly longer survival times than those infected with non-ST5 strains (P <0.0001, Mantel-Cox log rank test) (Panel A). Two ST5 strains, BK147 and BMD1646, were attenuated, suggesting high degree of heterogeneity within the ST5 cluster. Mice infected with BK147 and BMD1646 survived for as long as 42 days post-infection, at which point the experiment was terminated and all infected mice sacrificed (Panel B).

The tissue-specific fungal burden data are shown in Table 3. All organisms established lung infection and disseminated brain infection as early as 7 days post infection. Non-ST5 infections resulted in higher fungal burdens in lung than ST5 infections at all time-points (day 7, *p*<0.001; day 14, *p*<0.0001; and end of the experiment, *p*<0.0001 (Figure 3). Again, these data were not driven by the two apparently attenuated ST5 isolates as the fungal loads associated with the ST5 infections were significantly lower even when excluded from the analysis. The majority of animals infected with the attenuated ST5 organisms (BMD1646 (4/5 mice) and BK147 (5/5 mice)) had low fungal burdens in the lungs up to day 14 (Figure 3). Fungal burdens in the brain were higher in non-ST5 infected animals at all time points; however, this difference was only statistically significant at the end of the experiment (*p*=0.054, *p*=0.36, and *p*=0.01; Mann-Whitney test at days 7, 14 and point of impending death (time when 15% body weight loss is observed).

**TABLE 3.** Tissue fungal burden in lung and brains of mice at days 7, 14 and at time of death by infecting MLST Sequence Type (ST)

| Fungal burden |  |  |  |
| --- | --- | --- | --- |
| (Mean log<sub>10</sub> colony forming units per gram of tissue) |  |  |  |
| [95%CI] |  |  |  |
| Tissue | ST5 | Non-ST5 | P Value |
| Lung (Day 7) | 4.96 | 7.07 | <0.001 |
|  | [3.86 - 6.05] | [6.85 - 7.29]* |  |
| Brain (Day 7) | 0.81 | 1.59 | 0.054 |
|  | [0.38 - 1.24] | [0.87 - 2.30] |  |
| Lung (Day 14) | 5.48 | 8.00 | <0.0001 |
|  | [4.34 - 6.62] | [7.77 - 8.23]* |  |
| Brain (Day 14) | 1.56 | 1.91 | 0.36 |
|  | [0.81 - 2.31] | [1.12 - 2.69] |  |
| Lung (Death) | 3.81 | 6.53 | <0.0001 |
|  | [3.05 - 4.57] | [6.30 - 6.76]* |  |
| Brain (Death) | 2.90 | 4.29 | 0.01 |
|  | [2.23 - 3.56] | [3.90 - 4.68]* |  |

\*Mann-Whitney test.

**Figure 3.**
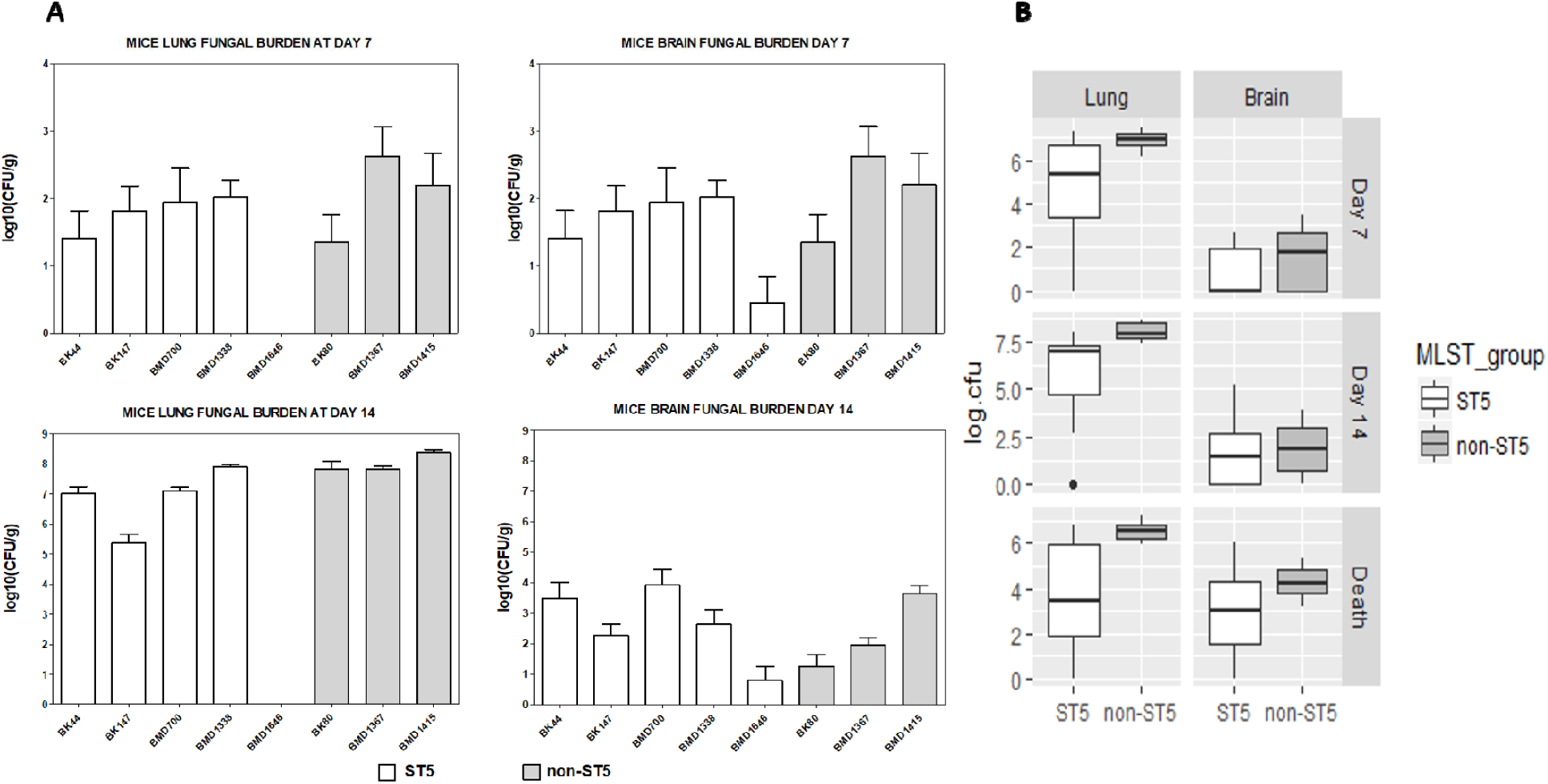
Fungal burden in mouse lung and brain tissue at days 7, 14 and the point of impending death (mortality experiment) according to infecting genotype. *In vivo* virulence (tissue fungal burden and mortality) was assessed in three independent mouse infection experiments. In the first two experiments, five A/J mice were infected with each of the eight test *C. neoformans* isolates (five ST5 isolates and 3 non-ST5 isolates, total N=40 mice in each experiment). All mice were sacrificed at day 7 post-infection in the first experiment and at day 14 post-infection in the second experiment to assess *in vivo* fungal burden in mouse lung and brain. Fungal burden in the lung and brain at either day 7 or day 14 for each isolate is presented in panel A. In the last experiment, ten A/J mice were infected with each of the 8 test *C. neoformans* isolate (five ST5 isolates and 3 non-ST5 isolates, N=80 mice in total) and monitored daily until the point when 15% of body weight loss, a sign of distress and impending death, was evident. Fungal burden in lung and brain at point of death was again assessed as before. Here, for each strain, five mice were randomly selected from the ten test mice for fungal burden investigation. The pooled fungal burden in non-ST5 infections was higher than in ST5 infections in both lung tissue at all time points, and in brain tissue at the point of sacrifice (Panel B) (For P values see text). Boxplots (Tukey’s method) describe the median and interquartile range, the whiskers demarcate the largest or smallest values that were not outliers (black dots); outliers are defined as more than 1.5 times the interquartile range from the nearest quartile.

### ST5 C. neoformans isolates induce of TNF-α in the lungs of infected mice

We next measured the cytokine concentrations in lung homogenates at days 7 and 14 post-infection (Figure 4). The concentrations of TNF-α were significantly higher in lung homogenates from mice infected with ST5 than non-ST5 infections (*p*=0.01). Of note, the highest day 7 lung TNF-α concentrations were associated with the ST5 organisms that were most attenuated in the mouse survival model (BK147 and BMD1646). Correspondingly, the lowest TNF-α concentrations were measured in the most virulent ST5 isolate in the mouse model (BMD1338). By day 14, the mean TNF-α concentrations in lung homogenates associated with the ST5 infections had declined from 3933.71 pg/g lung to 2802.36 pg/g lung (*p*<0.001). Other Th-1 cytokines including IL-12, IL-17, and IFN-γ also decreased in ST5-infected mice between days 7 and 14 (*p*<0.001, *p*<0.01 and *p*=0.02, respectively). This contrasted with the non-ST5 infections, in which the mean TNF-α levels had increased from 3256.25 pg/g to 6378.76 pg/g (*p*=0.02) over this same period. The high TNF-α concentration associated with non-ST5 infections at day 14 was driven by a single isolate - BMD1415. The TNF-α concentrations in in lung homogenates from mice infected with this isolate were significantly higher than for infections with any other isolate of any lineage (*p*<0.001). There was no statistically significant differences in TNF-α concentrations at day 14 between lineages when this isolate was excluded from the analysis. Similarly, the concentrations of these two cytokines in BMD1415-infected mice was significantly higher than in mice infected with any other isolate (*p*<0.001). However, the IL-12 concentrations declined in non-ST5-infected mice, including BMD1415, between day 7 and 14 (4 fold decrease, *p*<0.0001) (Figure 5).

**Figure 4.**
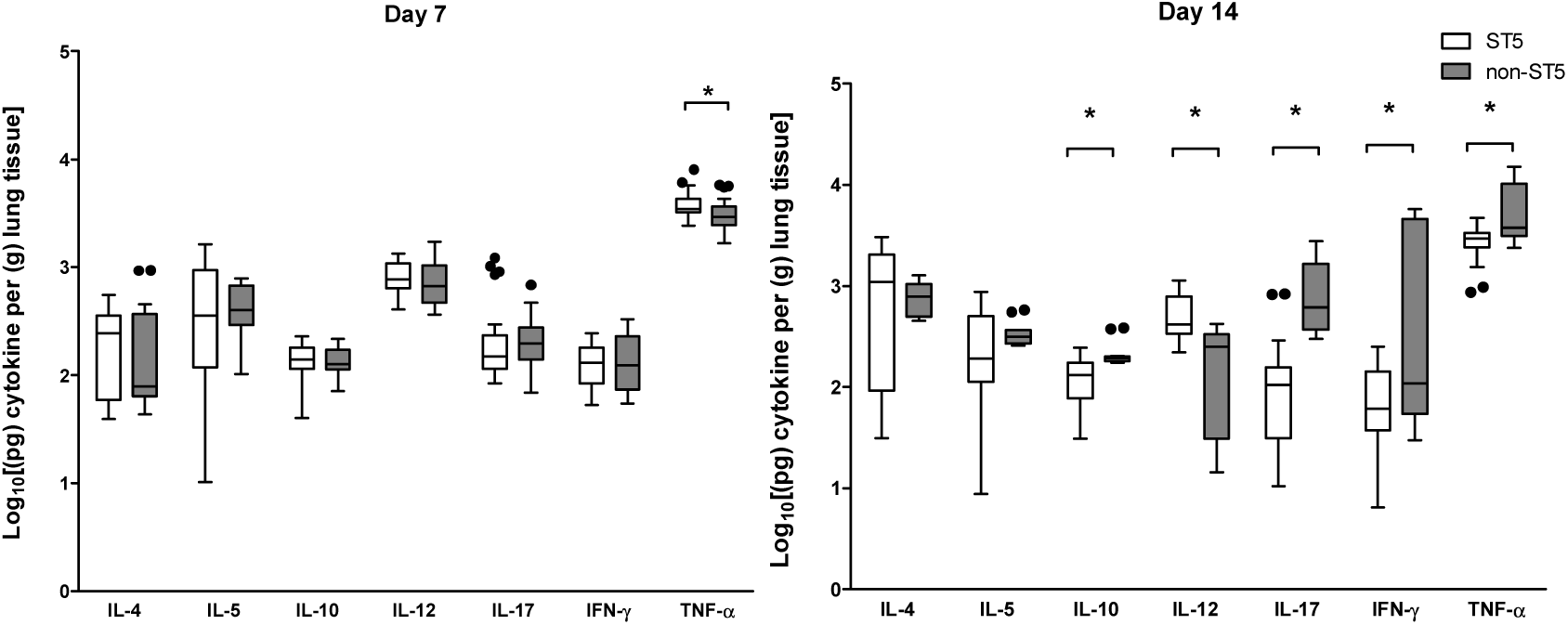
Genotype-specific cytokine concentrations from lung homogenate of A/J mice at 7 and 14 days post infection with 5×10^4^ *C. neoformans* cells/mouse. Five mice were infected with each strain from each genotype at each time points. ST5 strains induced significantly higher levels of TNF-α at day 7, suggesting an earlier and more profound initial inflammatory response in infected mice. By day 14 mice infected with non-ST5 strains have higher levels proinflammatory cytokines, probably a result of ST5 yeasts being cleared more rapidly from infected mice. The horizontal line within the box indicates the median; boundaries of the box indicate the 25^th^ and 75^th^ percentile and the whiskers indicate the highest and lowest values of the results; outliers are denoted as black dots (Tukey’s method). Data are standardized as picograms of cytokine per gram lung tissue. Asterisks indicate statistically significant differences (Mann-Whitney test).

**Figure 5.**
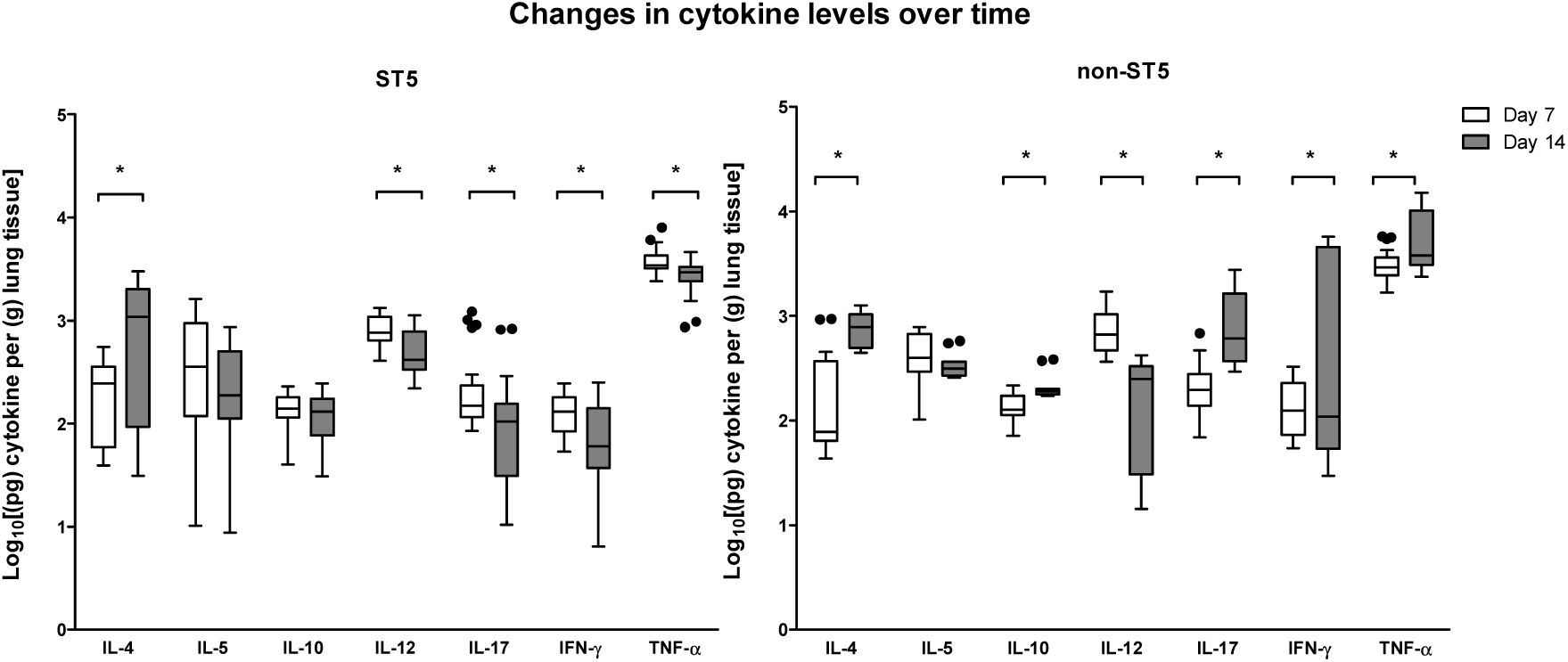
Genotype-specific changes in cytokine concentrations from lung homogenate of A/J mice infected with *C. neoformans* between day 7 and day 14 post-infection. Box and whisker plots (Tukey’s method) compare levels of each cytokine between day 7 and day 14 for each genotype. Data are standardized as picograms of cytokine per gram lung tissue. Asterisks indicate statistically significant differences (Mann-Whitney test).

We measured the TNF-α:IL-10 ratio as a proxy for the relationship between a Th-1 and a Th-2 response ^25^. From day 7 to day 14 days post-infection, the TNF-α:IL-10 ratio decreased by a factor of 0.78 in mice infected with ST5 isolates, while those infected with non-ST5 strains exhibited an increase in the ratio of 1.25-fold. However, when BMD1415 (ST4) was excluded from the analysis the TNF-α/IL-10 ratio in non-ST5 infected mice decreased by a factor of 0.67. By day 14 we detected elevated concentrations of IL-17 and IFN-γ in non-ST5 infected mice (5-fold and 17-fold increments, respectively (Table 4), but not in ST5 infected mice. Again, this effect was largely associated with BMD1415, since the concentrations of IL-17 and IFN-γ induced by this isolate at day 14 were significantly higher than for all others (*p*<0.001).

### Histopathological examination

Histological examination of infected lung tissue revealed evidence of inflammation, hemorrhage, edema and necrosis in most cases. These changes were generally greater by day 14 in comparison to day 7. There were no clear differences in histological scores between ST, other than with BMD1646 (ST5) which generated only mild inflammation with no evidence of necrosis or hemorrhage (Figure S5). PAS staining revealed extensive perivascular infiltration of leukocytes in mice tissue associated with both infecting genotypes (BMD1338-ST5 and BMD1415-ST4, Figure 6). Extensive *in vivo* encapsulation of *C. neoformans* var. *grubii* in mouse lung was visualized using Mucicarmine staining (Figure 7). However, it was unknown whether the bigger capsule and cell size associated with ST5 organisms *in vitro*, as described earlier, was also evident *in vivo* since we were unable to perform this measurement.

**Figure 6.**
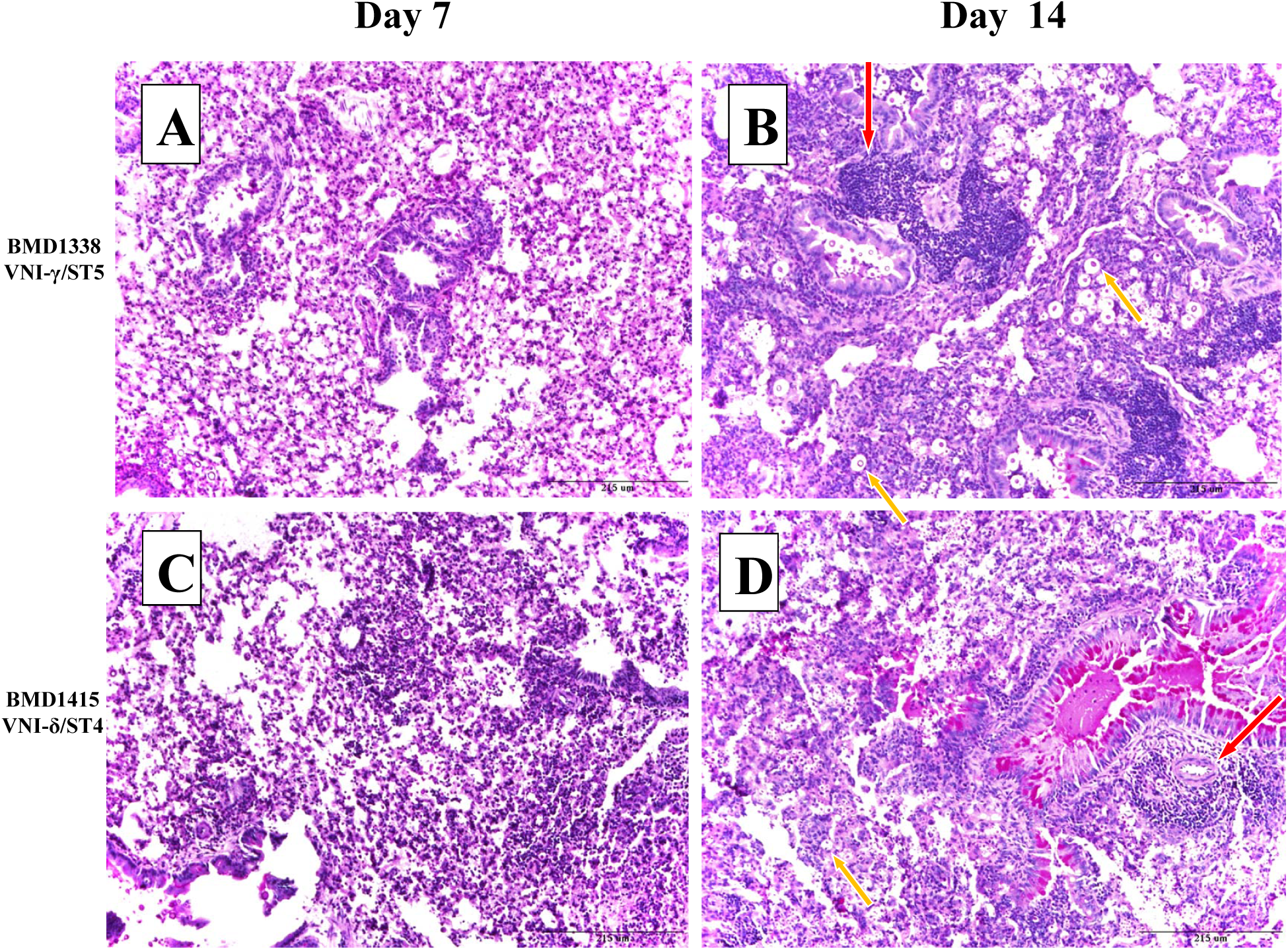
Periodic Acid Schiff (PAS) staining of pulmonary tissue from mice infected with BMD1338 and BMD1415 on days 7 and 14. The two strains represent ST5 and non-ST5, respectively. A/J mice were inoculated intranasally with 5×10^4^ yeast cells. Lung specimens were harvested at days 7 and 14 for histopathological examination. Photomicrographs were obtained at 200X magnification; the scale bar represents 215μm). (A-B): lung sections from mice infected with BMD1338 (VNI-γ/ST5) at day 7 and day 14, respectively. (C-D): lung sections from mice infected with BMD1415 (VNI-δ/ST4) at day 7 and day 14, respectively. Perivascular infiltration (red arrows) and necrosis are more marked by day 14 for both strains. Encapsulated yeasts (yellow arrows), notable for the larger cell size and capsule thickness of BMD 1338 compared with BMD1415.

**Figure 7.**
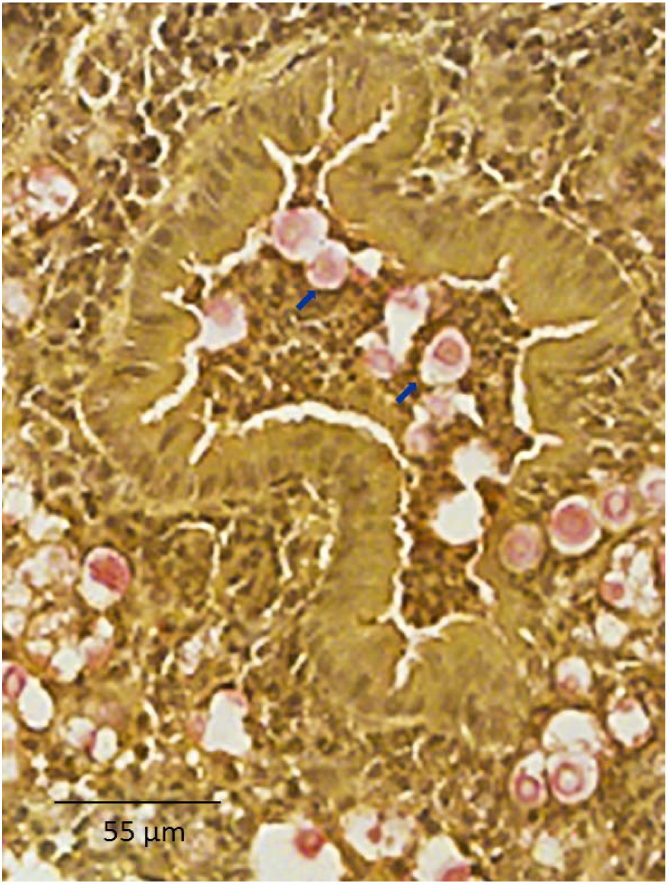
Mucicarmine staining of capsular material in paraffin-embedded mice pulmonary tissue. Uninflated lung specimens were harvested from mice as described in the methods. Mucicarmine staining was performed to visualize the cryptococcal capsule. Photomicrographs were captured at 400X magnification with scale bar indicating 55 µm. Capsular polysaccharide is stained pink (indicated by blue arrows), demonstrating diffuse localization consistent with extensive capsule production by yeasts in the alveolar space.

## Discussion

Most cases of cryptococcal meningitis in HIV uninfected, apparently immunocompetent, patients in Vietnam and East Asia are due to *C. neoformans* strains of multi-locus sequence type 5 (ST5) ^9,10,12,17,26–28^. This phenomenon could be explained either by ST5 strains being intrinsically more pathogenic, or due to unidentified lineage–specific host immune defects or exposures. It is unlikely that the high prevalence of ST5 infection observed in HIV-uninfected patients is explained by a significantly greater prevalence of the lineage in the environment since it causes only 35% of cases in HIV-infected patients within the same geographical area ^9^. Furthermore, data from China suggest that ST5 strains are significantly less prevalent in the environment, making up only 5% of isolates recovered in a recent study ^29^. We investigated the first hypothesis by comparing previously identified *in vitro* virulence-associated phenotypes, along with murine *in vivo* virulence and immune responses, between lineages. All isolates were derived from HIV infected or uninfected Vietnamese patients with cryptococcal meningitis. The comparison by MLST defined lineage is appropriate because ST5 is a coherent and distinct group, whole genome sequence data revealing that there are few intra-lineage genomic variations between ST5 strains ^17^. More recent phylogenetic analysis using whole genome sequencing of a larger collection of clinical isolates from Vietnam, including the eight strains tested in the mouse model, has established that the MLST ST5 and ST4/ST6/306 groups are genetically distinct lineages (VNIa-5 and VNIa-4, respectively) ^15^. It is unlikely that any phenotypic variations among ST5 isolates are primarily attributable to *subtle* intra-lineage genomic variations since phenotypic and genotypic diversity are not tightly coupled in *C. neoformans* var. *grubii* ^30^. Due to the fact that ST5 has been shown to be consistently associated with the clinical phenotype of interest (infection in HIV-uninfected patients), and that strains from HIV uninfected patients are dispersed throughout the VNIa-5 cluster, we believe the ability to infect apparently immunocompetent hosts is common to all ST5 isolates. All comparisons of ST versus non-ST5 in this study were thus essentially VNIa-5 versus VNIa-4. We found that isolates from all STs were able to grow in *ex vivo* human CSF and at 37°C - essential characteristics for establishing human CNS infection. While these qualities would be needed for disease in both HIV infected and immunocompetent patients, it might have been expected that ST5 strains would grow more rapidly in these conditions. The lack of ST-specific differences in these phenotypes suggests that the ability to establish disease in HIV-uninfected/immunocompetent patients is not driven by simple adaptations to these conditions. Noticeably, ST5 cells were significantly larger than non-ST5 cells, had thicker capsules *in vitro*, and had more within lineage variation in these characteristics. Capsule size and composition are known to vary during infection and under specific stress conditions ^31,32^, influencing macrophage phagocytosis and modulating host immune response 32–35, and in human disease, *ex vivo* capsule size has been associated with higher intracranial pressures, slower yeast clearance and attenuated inflammation ^34^.While we did not formally measure yeast cell or capsule size in our *in vivo* experiments, mucicarmine staining was suggestive that capsules were indeed larger during mouse infection. It is possible that the ability of ST5 strains to cause infections in immunocompetent patients is a function of increased responsiveness to capsule-inducing conditions. Further investigation of genotype-specific characteristics of *in vitro* and *in vivo* capsular polysaccharide production, composition, and morphology, may elucidate a specific role in ST5-associated pathogenesis. Of note, cryptococcal virulence factors frequently have additional metabolic functions; the increased cell and capsule size seen in ST5 isolates may be a side effect of other processes involving capsular-biosynthesis genes (for example carbon source sensing, sugar transport and spore formation) ^36,37^.

Our data indicate that phenotypic heterogeneity may be a hallmark characteristic of the ST5 lineage. Heterogeneity is a desirable trait for microbial populations under selection pressure allowing the exploitation of, and survival, in novel niches ^38^. Chow et al (2008) has previously reported that ST5 *C. neoformans* var. *grubii* possesses unique genomic features which may drive niche adaptation ^39^. We speculate that the phenotypic heterogeneity associated with strains from the ST5 lineage is a strategy that facilitates successful colonization of novel environmental niches, including the exploitation of infrequent specific human immune deficits. Morphological variation including cell size, capsule size and cell shape has also been associated with different patient clinical symptoms, suggesting greater capacity for pathogenicity, immune evasion and pathogenesis ^40^.

However, paradoxically, we found no evidence that ST5 isolates have greater virulence in the mouse model. There are several possible explanations. First, virulence in mice is variable depending on mouse breed and may not accurately reflect the immunological heterogeneity of the human population ^41^. Second, yeasts with different pathogenic potentials, associated with their isolation from different sources (i.e. clinical versus environmental, immunocompromised versus immunocompetent patients) may have the same or paradoxical pathogenic potential in experimental animal models. An example is *Cryptococcus gattii* which is associated with infection in immunocompetent patients and therefore is considered to be more fit in the human host than *C. neoformans*. However, the hypervirulent *C. gattii* strain R265, responsible for the on-going Vancouver outbreak, has similar virulence in both C57BL/6 and A/J mice to the *C. neoformans* H99 strain, which was derived from a patient with Hodgkin’s disease on chemotherapy ^42^. Third, the A/J mouse breed is not immunologically intact; it may be an imperfect model of infection for immunocompetent hosts ^43^. Rather, A/J mice may be a better model of disease in immunosuppressed patients, as they are highly susceptible to cryptococcal disease, and the patterns of cytokine expression in mice with disseminated cryptococcosis are similar to those seen in HIV-infected patients with CM ^44^. Consistent with this, we could not detect clinical differences in disease course or outcome between HIV patients infected with ST5 versus other strains in Vietnam ^17^. Models that better mimic infection in immunocompetent hosts are needed.

We did identify lineage specific differences in immune response in the mouse model. Previous research has suggested that a Th1 type immune response, defined by the TNF-α/IL-10 ratio, is protective, and a Th2 response is associated with poor outcomes ^45,46^. We found no evidence of genotype-specific differences in TNF-α/IL-10 ratios by lineage in the murine infection model. Rather, we found higher initial (day 7) TNF-α concentrations in mice infected with ST5 isolates, suggesting this genotype elicits a more intense initial inflammatory response. Previous studies have suggested that capsule components, or cryptococcal cells themselves, have a dose-dependent ability to stimulate TNF-α production by various immune effector cells ^47,48^. The more robust initial inflammatory response we observed may have been due to the ST5 capsular phenotype. Previously, it has been suggested that the ability of *C. gattii* to cause disease in apparently immunocompetent patients is because it induces a less severe inflammatory response compared with other cryptococcal species ^42^. The robust initial inflammatory responses seen in our murine infection experiments are not consistent with this being the mechanism underlying the ability of ST5 *C. neoformans* var. *grubii* organisms to cause disease in the immunocompetent.

*In vivo* controlled infection studies in mice, including ours, commonly employ the classic definition of pathogenicity as the microbe’s capability to cause disease in a susceptible host, whereas virulence corresponds to the severity of the ensuing pathology ^49^. Using the same infective dose for all strains we failed to demonstrate that ST5 strains had greater virulence.

The difference we observe in prevalence of different lineages in immunocompetent and immunosuppressed humans may actually represent specific differences in pathogenicity - the ability of the organisms to colonize the host and establish infection. We could not assess this with our experimental system.

In summary, our cohort of ST5 *Cryptococcus* isolates displayed two notable phenotypes. First, despite their well-documented ability to cause disease in HIV uninfected humans, they appeared to be less virulent in a murine model than the other sequence types, as demonstrated by reduced fungal burdens in tissue and prolonged mouse survival. Second, ST5 strains had larger capsules and cell sizes than the other genotypes, and greater variability in this phenotype throughout the lineage. These data lead us to the following conclusions. First, clinical isolates, which have by their nature already undergone selection within the human host, can possess wide variability in the expression of virulence phenotypes within a single lineage. Secondly, the use of host risk factors and immune phenotypes to derive an understanding of the factors that drive the pathogenicity of *Cryptococcus neoformans* may be more complex than anticipated. Associations may be difficult to make due to the relevance of the particular *in vitro* phenotypes, the animal models used, within strain heterogeneity, and population substructure. Moreover, there may be heterogeneity in the immune response of apparently immunocompetent patients which selects particular sub-cohorts of isolates of the same lineage. Laboratory phenotyping of larger numbers of clinical isolates is needed to define the lineage-specific differences that determine different human disease phenotypes.

Finally, it is possible that the categorization of strains into specific clades with limited genetic information such as MLST may lack precision to understand the relative fitness of specific strains in the human host. It is likely that whole genome sequencing will provide better mapping of the relationships between strains and virulence.

In conclusion, in this study, we demonstrated genotype-specific differences in *in vitro* and *in vivo* virulence phenotypes between *C. neoformans* var. *grubii* strains isolated from host with different immune status. However, there was also significant variation among strains isolated from apparently immunocompetent patients in specific *in vitro* and *in vivo* phenotypes tested. This higher rate of phenotypic variation may represent an evolutionary strategy for *C. neoformans* var. *grubii* to take advantage of novel niches and contribute to their ability to infect apparently immunocompetent hosts, despite generally being less virulent in a mammalian animal model. Furthering the understanding of the pathogenesis of cryptococcal meningitis will require investigation of large numbers of strains with associated robust clinical information, and the development of high throughput laboratory phenotypic studies that have clinical relevance in humans.

## Acknowledgements

This work was supported by a Wellcome Trust Intermediate Fellowship awarded to JND (WT097147MA), and a Henry Dale Fellowship jointly funded by the Wellcome Trust and the Royal Society (100087/Z/12/Z) awarded to SGB. Multiplex cytokine profiling was performed in the Immunology Unit of the Duke Regional Biocontainment Laboratory, which received partial support for construction from the US National Institutes of Health, National Institute of Allergy and Infectious Diseases (UC6-AI058607). The study was approved by the scientific committee of the Hospital for Tropical Diseases, Ho Chi Minh City, Vietnam.

## FIGURE LEGENDS

**FIGURE S1. Phenotyping of *C. neoformans* var. *grubii* isolates from Vietnam (ST5: n=15 and non-ST5: n=15).** Data were obtained from 30 isolates (15 ST5 and 15 non-ST5) in 3 different experiment batches, with 3 technical replicates of each isolate per batch. Data are expressed as the ratio between measurements of the test isolates and that of H99. Boxplots (Tukey’s method) describe the median and interquartile range (panel A-C). For assessment of urease activity (panel D), we used the time to complete color change of the agar plate as an indirect measure of extracellular urease activity since all *C. neoformans* var. *grubii* isolates in our collection were positive for urease activity. Each isolate were tested in triplicate and monitored in real time by live-imaging. No significant difference in any virulence-associated phenotype were observed between the two MLST groups.

**FIGURE S2. Growth at different temperature of 8 *C. neoformans* (five ST5 and three non-ST5) isolates from Vietnamese patients used in the murine infection experiments.** All strains expressed similar growth at 30°C. Two ST5 isolates (BMD1646 and BK147) displayed diminished growth at 37°C compared to other test isolates.

**FIGURE S3. *Ex vivo* growth in human cerebrospinal fluid (CSF) of 8 clinical *C. neoformans* isolates from Vietnamese patients that were tested for virulence in mice, representing both ST5 (n = 5) and non-ST5 (n = 3) genotypes.** The same inoculum of yeasts were inoculated in pooled human CSF and incubated at 37°C. The inoculated CSF was serially diluted and plated on YPD agar at day 1 and day 3 post-inoculation. The wild-type H99 strain and the H99-derived mutant *Δena1*, lacking a cation ATPase transporter which results in decreased viability in human CSF and within macrophages, were included as controls. An apparent difference in growth between ST5 and non-ST5 isolates could be observed at day 3.

**FIGURE S4. Melanin production on L-DOPA medium of eight clinical *C. neoformans* isolates from Vietnamese patients representing both S5 (n =5) and non-ST5 (n = 3) genotypes that were tested for virulence in mice.** For each isolates, 10μl of 10^6^ cell suspension was inoculated on L-DOPA agar and incubated in the dark at 30°C and 37°C. No clear genotype-specific patterns of melanization were observed at either 30°C or 37°C. The five ST5 strains displayed marked variation in the degree of pigment production, from mildly melanized (BK147) to highly melanized (BK44 and BMD1646).

**FIGURE S5: Histopathological scores across 4 categories of tissue damage (Inflammation, necrosis, hemorrhage and edema).** Thin sections of paraffin-embedded lung specimens from infected mice were stained using the Periodic Acid Schiff (PAS) method. Specimens were assessed by an independent pathologist who was blind to infecting isolate and in randomized order. Scores ranged from 0 (no changes) to 10 (severe changes), corresponding to the severity of pathology in in each category as per the Duke Veterinary Diagnostic Laboratory protocol (Division of Laboratory Animal Resources).

