## Supplementary figures and images for "Assessing the virulence of *Cryptococcus neoformans* causing meningitis in HIV infected and uninfected patients in Vietnam"

### Supplemental Figure S1

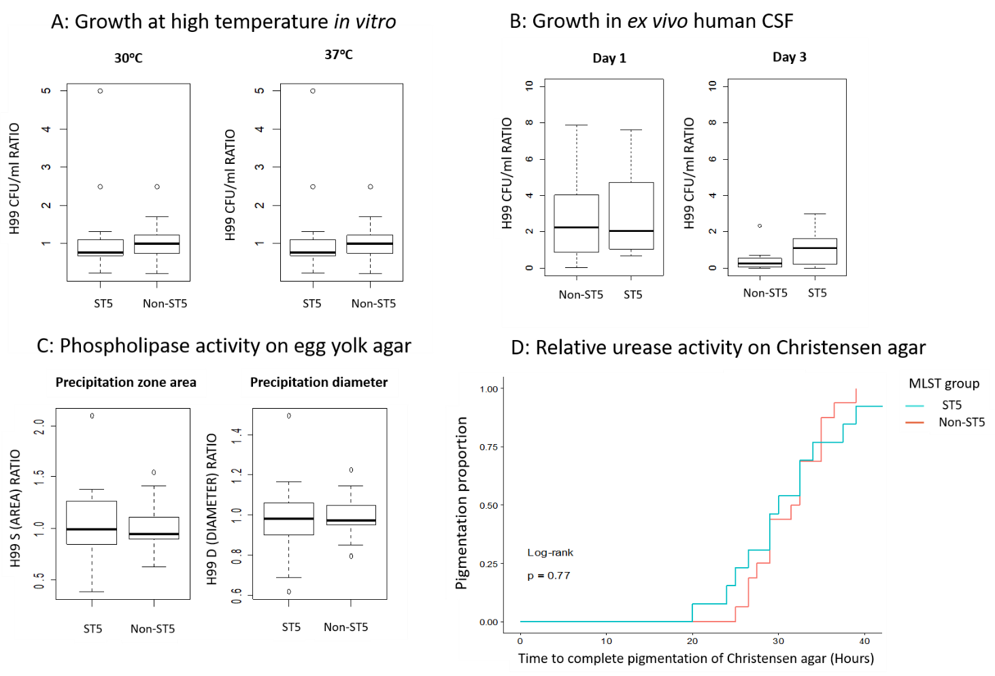

### Supplemental Figure S2

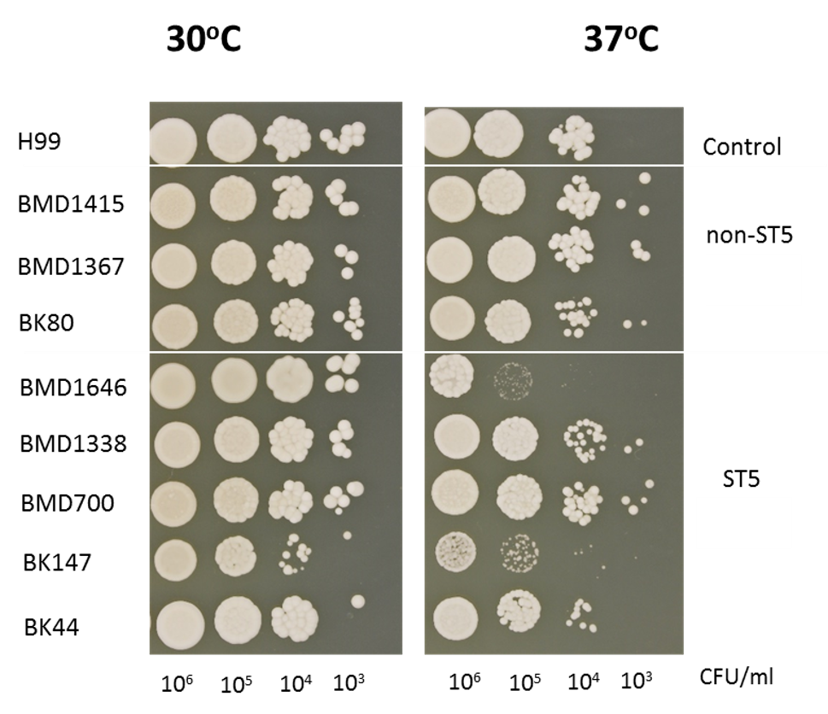

### Supplemental Figure S3

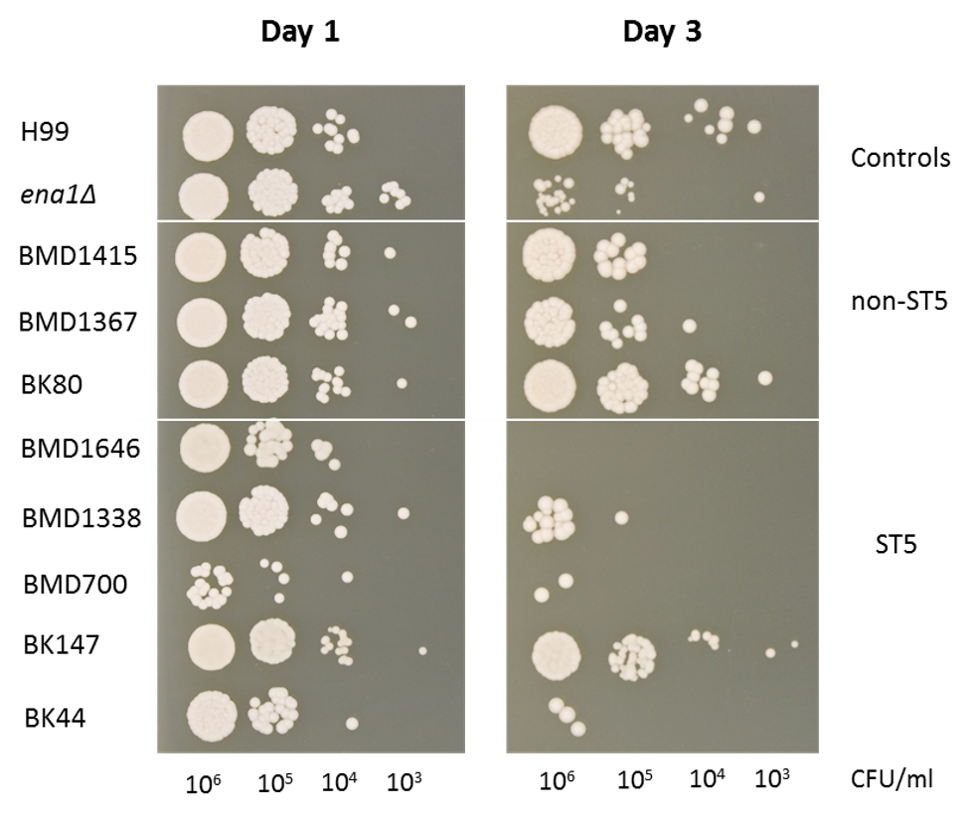

### Supplemental Figure S4

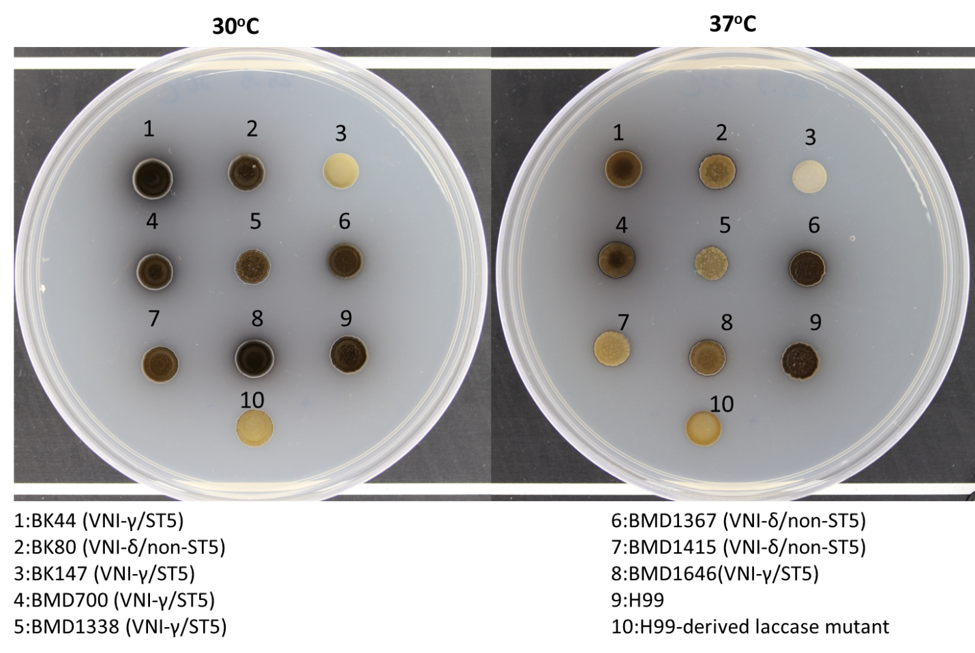

### Supplemental Figure S5

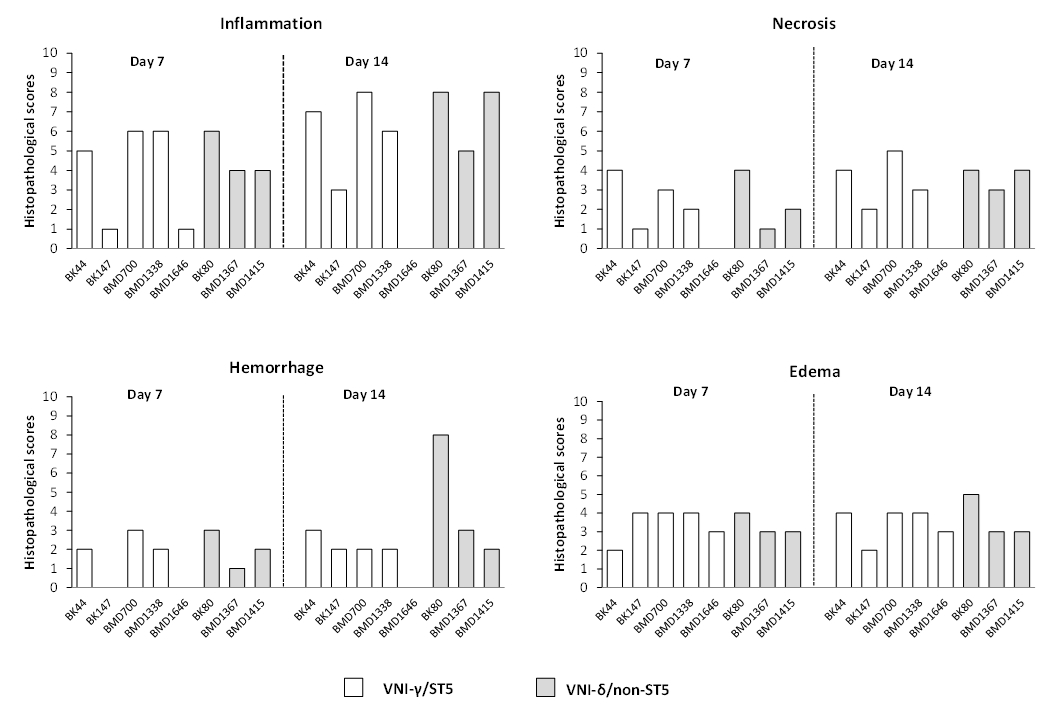
